# Synthetic diet feeding exacerbates central nervous system autoimmunity in mice independent of weight gain

**DOI:** 10.64898/2026.11.13.766400

**Authors:** Alesja Dernst, Benedikt Gansen, Johan B. Kantsjö, Alessio Murgia, Arijan Pasic, Svitlana Rozanova, Frederik Heinrich, Pawel Durek, Titus Schlüter, Yubell Alvarez, Romina Kaiser, Maia Cristodaro, Dennis Marinus De Graaf, Lukas Jacobi, Dariya Rainio, Bernardo S. Franklin, Mir-Farzin Mashreghi, Katalin Barkovits, Katrin Marcus, Federica Sallusto, Eicke Latz, Francesca Ronchi, Felix Meissner, Felix D. Weiss

## Abstract

Consumption of a nutrient deficient diet high in saturated fats and sugar is associated with the increased incidence and exacerbation of inflammatory and autoimmune diseases. Efforts to resolve the molecular mechanisms underlying diet-dependent changes to immune responses rely on the use of synthetic diets in mice, often using a standard laboratory chow as control. Critically, it remains unknown whether synthetic diet specific alterations in nutrient and metabolite availability, independent of weight gain and exposure to dietary fats and sugar, can affect the immune response. We reveal that exposure to a synthetic diet, independent of weight gain, is sufficient to alter the immune response, driving increased severity in the experimental autoimmune encephalomyelitis (EAE) model of central nervous system (CNS) autoimmunity. Mechanistically, synthetic diet feeding leads to systemic reductions in short-chain fatty acid (SCFA) abundance, and supplementation of synthetic diet with butyrate mitigates the excessive production of pathogenic T cells, and ameliorates synthetic diet driven increases in EAE severity. Finally, we identify the presence of peripheral Tregs (pT_regs_) in the CNS of EAE mice, which can arise from shared clonal ancestors as T helper (Th) cells, and reveal that butyrate supplementation increases the proportion pT_regs_ to Th17 cells during EAE.

## Introduction

Differences in dietary habits, particularly the consumption of a Western diet high in fat, sugar and cholesterol, are known to drive the development of non-communicable diseases such as atherosclerosis and type-II diabetes, and have also been implicated in the onset or exacerbation of neurodegenerative^1,2^ and autoimmune diseases, including multiple sclerosis^6–9^. Mouse models of human dietary habits have expanded our understanding of the molecular mechanisms that underpin the relationship between differential dietary exposure and disease development, with diet induced changes to metabolite availability, lipid abundance and microbial composition all shown to alter both the innate and adaptive immune compartments^8–11^.

Globally, laboratory mice are maintained on “chow” diets largely derived from natural grains such as wheat and corn, and supplemented with additional vitamins and trace elements. Mouse models of human diets on the other hand are synthetically made, where nutrients are provided by purified ingredients, with significant variations in macronutrient sources, particularly dietary fibres, relative to normal chow^12–14^. While synthetic high-fat and Western diets have been shown to affect disease development, often in comparison to chow controls^15^, there remains a limited understanding of whether synthetic diet exposure alone, independent of weight gain or exposure to increased levels of dietary fats and sugars, reprograms the immune compartment. In this study we revisit the assumption that synthetic diets serve as neutral experimental controls, and analyse the effect of synthetic diet feeding on the EAE mouse model of CNS autoimmunity, which was previously shown to be exacerbated by exposure to a synthetic high-fat diet relative to chow^16–19^.

We find that synthetic diet feeding alone, independent of weight gain or increased exposure to dietary fats and sugar, leads to systemic reductions in dietary fibre metabolites, promotes the formation of pathogenic Th17 cells, and drives increased clinical severity in EAE. These findings demonstrate that synthetic diets, even in the absence of increased dietary fats and sugars, are pro-inflammatory and can augment disease development in murine models.

## Results

### Synthetic diet feeding drives increased clinical severity and demyelination in the EAE mouse model of CNS autoimmunity

In order to determine whether synthetic diet feeding alters the severity of CNS autoimmunity, we fed 8-week-old C57B6/J mice a synthetic Western diet (high in fat, sugar and cholesterol), a synthetic control diet, or standard laboratory chow, for a period of 8 weeks prior to, and throughout, EAE induction (**Figure 1A**). As expected, Western diet fed mice gained significantly more weight than mice fed either synthetic control diet or chow (**Figure 1B, C**). Strikingly, however, synthetic control diet fed mice experienced the greatest increase in EAE severity relative to chow (**Figure 1D**). In addition, spinal cords of synthetic control diet fed mice displayed a significant increase in the number of demyelinated lesions during EAE (**Figure 1E, F**), as well as a significant increase in lesion size (**Figure 1G**), relative to chow. We therefore focused on resolving the mechanisms that drive increased disease severity in synthetic control diet fed mice relative to chow.

**Figure 1.**
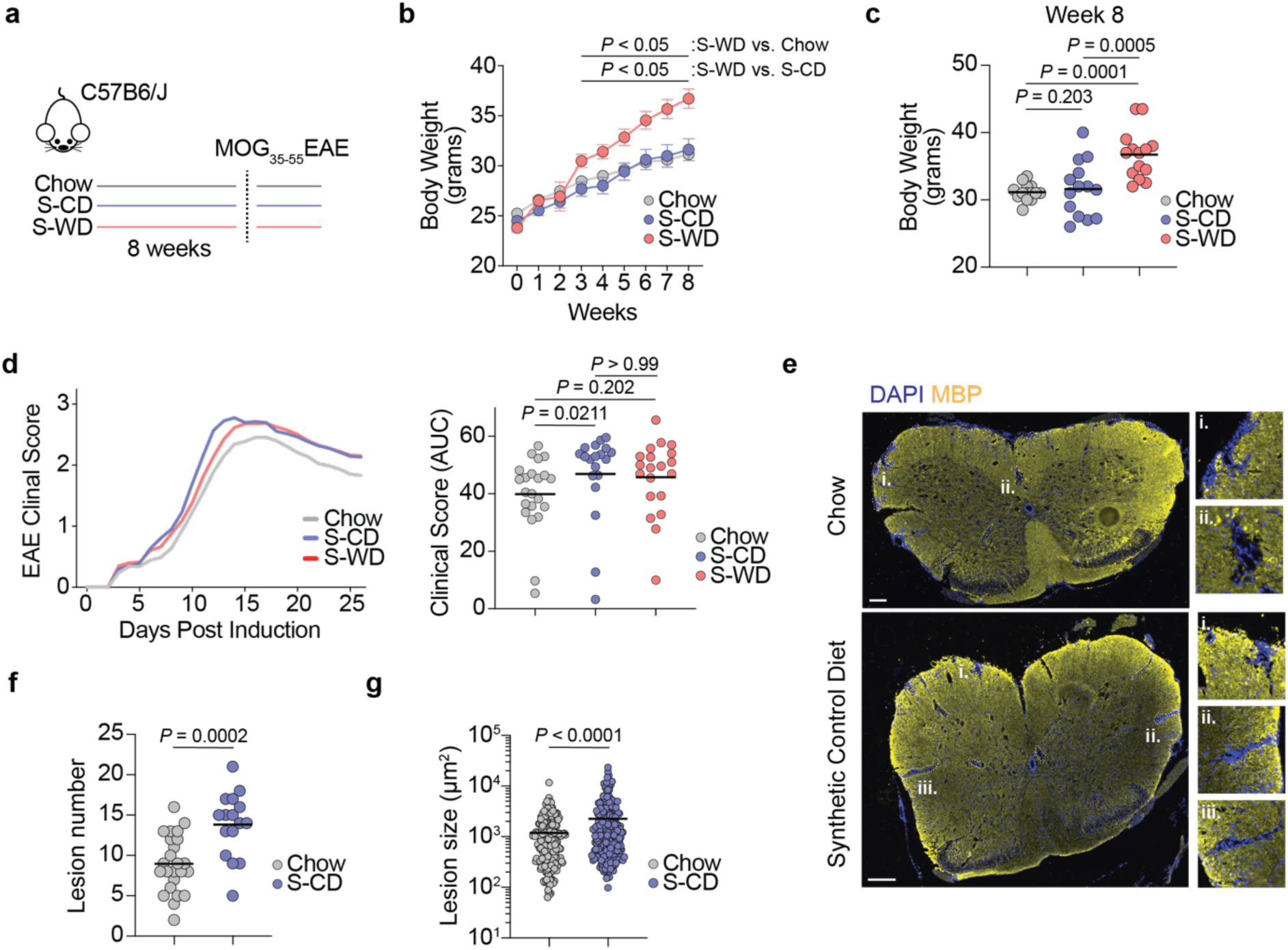
Synthetic diet feeding exacerbates clinical severity in the EAE mouse model of neuroinflammation. **a.** Schematic representation of feeding and EAE induction paradigm (S-CD = synthetic control diet; S-WD = synthetic western diet). **b.** Body weight in grams of mice over 8 weeks of chow, S-CD or S-WD feeding. Mean ± s.e.m, *P* values calculated using two-way repeated measures ANOVA with Tukey’s multiple comparisons test, (n = 14 per group). **c.** Body weight in grams of mice following 8 weeks of chow, CD or WD feeding. Black bars indicate mean values. *P* values calculated using ordinary one-way ANOVA with Tukey’s multiple comparison test, (n = 14 per group). **d.** Left: EAE clinical severity score over time; Right: AUC (area under curve) analysis of EAE clinical scores from chow, S-CD and S-WD fed mice. Black bars indicate mean values. *P* values calculated using Kruskal-Wallis test and Dunn’s multiple comparisons test. (n = 22 (Chow), n = 19 (S-CD), n = 20 (S-WD); data from two independent experiments). **e.** Representative immunofluorescence images of EAE lesions from chow and synthetic control diet fed mice labelled for DAPI and myelin basic protein (MBP). Lesions are highlighted with white dashed lines. Scale bars indicate 100µm. **f.** Number of lesions per spinal cord section in chow and synthetic control diet (S-CD) fed mice. Bars indicate mean values. *P* value calculated using parametric two-tailed unpaired T-test. (n = 24 (Chow), n = 16 (S-CD); data from three biological replicates per condition). **g.** Lesion size in chow and synthetic control diet (S-CD) fed mice. Bars indicate mean values. *P* value calculated using Mann Whitney U test. (n = 215 (Chow), n = 221 (S-CD); data from three biological replicates per condition).

### Synthetic diet feeding promotes the induction of pathogenic Th17 cells and the recruitment of peripheral myeloid cells to the CNS during EAE

EAE is induced through the formation of pathogenic autoreactive RORγt^+^ Th17 cells^20–22^, with CNS inflammation and neurodegeneration further driven by the reprogramming of CNS-resident glia, including microglia^23–25^ and astrocytes^26^, and the recruitment of peripheral neutrophils and monocytes^27–34^. Synthetic diet feeding did not significantly alter the proteomes of FACS isolated CNS mononuclear phagocytes, Ly6C^hi^ monocytes or astrocytes at the peak of EAE relative to chow (**Supplemental Figure 1A, B**). Synthetic diet feeding did not affect the relative abundance of spinal cord CD80^+^, CD86^+^ or CD11c^+^ microglia and monocyte-derived cells during EAE (**Supplemental Figure 1C, D**), corroborating proteome data.

Synthetic diet fed mice however did display a significant increase in the proportion of spinal cord RORγt^+^ Th17 cells, with a concomitant significant reduction in the proportion of anti-inflammatory regulatory T cells (T_regs_; **Figure 2B**). In addition, synthetic diet feeding increased the relative abundance of T-bet^+^ and RORγt^+^ T-bet^+^ cells among all CD4^+^ T cells (**Figure 2C**). T-bet expression has previously been associated with pathogenic Th17 cells in EAE^35,36^, as well as the production of GM-CSF^37,38^. FACS analysis of cytokine production in spinal cord T cells isolated from synthetic diet and chow fed mice revealed a significant increase in the proportion of GM-CSF^+^ CD4^+^ T cells at EAE peak, with a similar trend in IL-17A^+^ (**Figure 2D, E**), while the proportion of IFNγ^+^ cells remained unchanged (**Figure 2E**).

**Figure 2.**
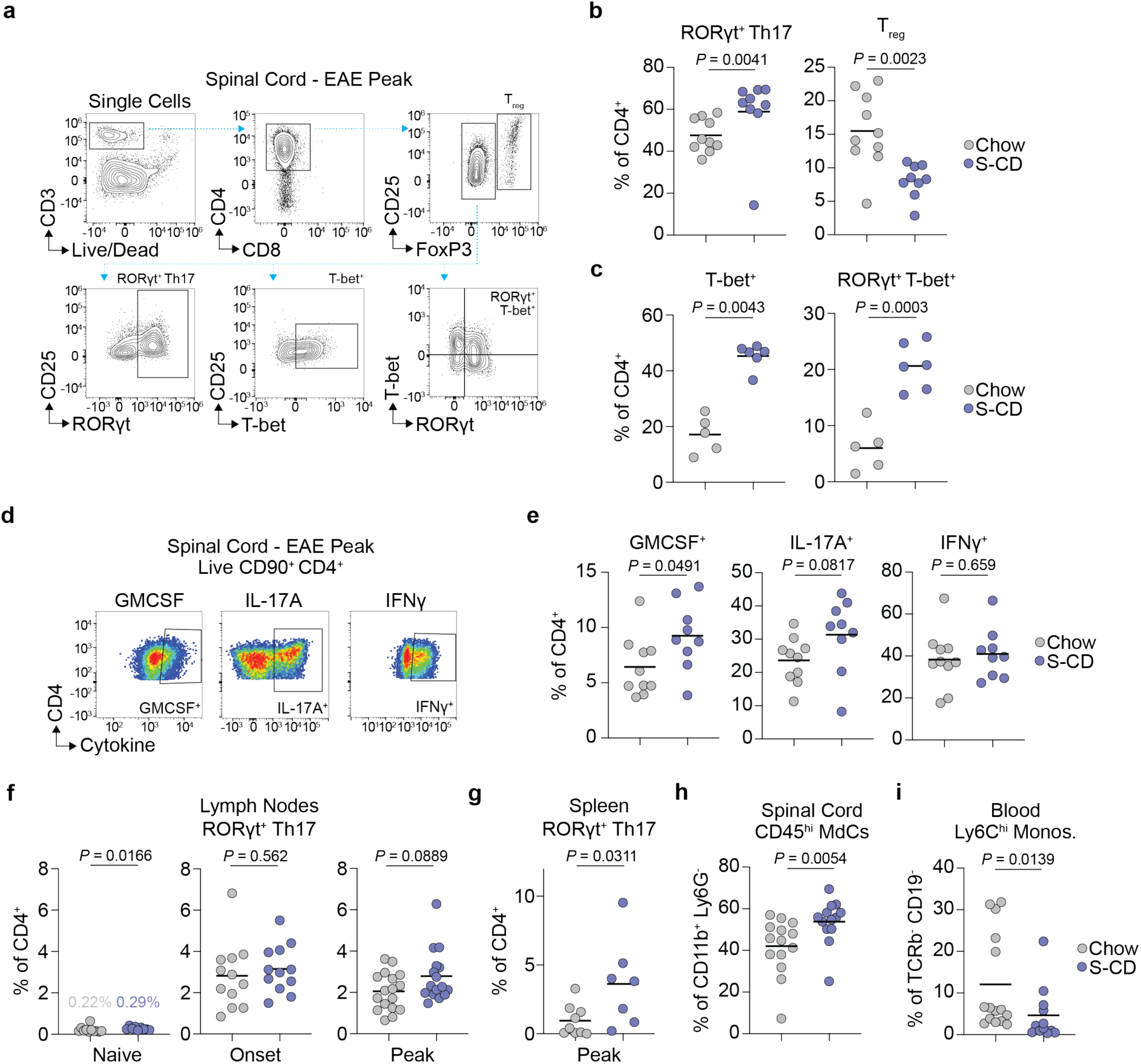
Synthetic diet feeding enhances spinal cord Th17 cell abundance and recruitment of peripheral monocytes during EAE. **a.** FACS gating strategy for the identification of RORγt^+^ Th17 cells, T_regs_, T-bet^+^ and RORγt^+^ T-bet^+^ cells from mouse spinal cord during EAE. **b.** Left: Proportion of RORγt^+^ Th17 cells from all CD4^+^ T cells in the spinal cords of chow and synthetic control diet (S-CD) fed mice at the peak of EAE. Right: Proportion of T_regs_ from all CD4^+^ T cells in the spinal cords of chow and synthetic control (S-CD) fed mice at the peak of EAE. Black bars indicate mean values, *P* values calculated using Mann-Whitney U test (RORγt^+^ Th17 cells) and Welch’s T-test (T_regs_). (n = 10 (Chow), n = 9 (S-CD); data from two independent experiments). **c.** Left: Proportion of T-bet^+^ cells from all CD4^+^ T cells in the spinal cords of chow and synthetic control diet (S-CD) fed mice. Right: Proportion of RORγt^+^ T-bet^+^ cells from all CD4^+^ T cells in the spinal cords of chow and synthetic control diet (S-CD) fed mice. Black bars indicate mean values, *P* values calculated using Mann-Whitney U test (T-bet^+^) and parametric two-tailed unpaired T-test. (RORγt^+^ T-bet^+^). (n = 5 (Chow), n = 6 (S-CD). **d.** FACS gating strategy of cytokine staining following restimulation (20nM PMA, 1µM Ionomycin, 10µg/ml Brefeldin A, 4 hours) of primary CD4^+^ T cells isolated from mouse spinal cords at the peak of EAE. **e.** Quantification of the % of GMCSF^+^, IL-17A^+^ and IFNγ^+^ CD4^+^ T cells from all CD4^+^ T cells isolated from the spinal cords of mice fed chow or S-CD during EAE following restimulation. Black bars indicate mean values, *P* values calculated using parametric two-tailed unpaired T-test. (n = 10 (Chow), n = 9 (S-CD); data from two independent experiments). **f.** Quantification of the % of RORγt^+^ Th17 cells from all CD4^+^ T cells in inguinal or CNS draining lymph nodes of naïve mice (left) and at EAE onset (middle) and EAE peak (right) from chow and S-CD fed mice. Black bars indicate mean values, *P* values calculated using Mann-Whitney U test (naïve & peak) and parametric two-tailed unpaired T-test (onset). (Naïve: n = 11 (Chow), n = 9 (S-CD); Onset: n = 12 (Chow), n = 12 (S-CD); Peak: n = 17 (Chow), n = 16 (S-CD), data from two independent experiments). **g.** Quantification of the % of RORγt^+^ Th17 cells from all CD4^+^ T cells in spleen at EAE peak from chow and S-CD fed mice. Black bars indicate mean values, *P* values calculated using Mann-Whitney U test. (n = 9 (chow); n = 7 (S-CD)). **h.** Quantification of the % of CD45^hi^ monocyte-derived cells (MdCs) among all mononuclear myeloid cells (CD11b^+^ Ly6G^-^) in the spinal cords of mice at EAE peak. Black bars indicate mean values, *P* values calculated using Mann-Whitney U test. (n = 14 (Chow), n = 13 (S-CD); data from two independent experiments). **i.** Quantification of the % of Ly6C^hi^ monocytes among all blood leukocytes (TCRb^-^ CD19^-^) at EAE peak. Black bars indicate mean values, *P* values calculated using Mann-Whitney U test. (n = 15 (Chow), n = 12 (S-CD); data from two independent experiments).

Initial T helper cell activation and polarisation occurs in the periphery during EAE^39,40^, followed by restimulation in the CNS. Prior to EAE induction, synthetic diet fed mice displayed a significant increase in the relative abundance of RORγt^+^ Th17 cells in inguinal lymph nodes compared to chow (**Supplemental Figure 2A; Figure 2F**). However, as expected, these comprised only a minor proportion of CD4^+^ T cells in the absence of immunisation (**Figure 2F**). Analysis of CNS draining lymph nodes during EAE revealed that the proportion of RORγt^+^ Th17 cells among all CD4^+^ T cells was unchanged at EAE onset in synthetic diet fed mice relative to chow (**Figure 2F**), had an upward trend at EAE peak (**Figure 2F**), and was significantly increased in spleen at EAE peak (**Figure 2G**). No differences were observed in the peripheral induction of Th17 cells between chow and synthetic diet fed mice following exposure to complete freund’s adjuvant in the absence of MOG_35-55_ peptide (**Supplemental Figure 2B**).

Th17 cell derived GM-CSF and IL-17A act to recruit peripheral monocytes to the spinal cord during EAE^31–34^, where they differentiate into monocyte-derived cells (MdCs), including macrophages and dendritic cells, and drive disease pathology. Synthetic diet fed mice had a significant increase in the proportion of CD45^hi^ MdCs relative to all mononuclear myeloid cells (CD11b^+^ Ly6G^-^) in the spinal cord at peak EAE relative to chow (**Figure 2H**; **Supplemental Figure 2C**). Consistent with increased monocyte extravasation, synthetic diet fed mice had a significant reduction in the proportion of Ly6C^hi^ pro-inflammatory monocytes among all circulating myeloid cells (TCRb^-^ CD19^-^; **Figure 2I; Supplemental Figure 2D**), despite showing no changes in bone marrow myelopoiesis prior to and during EAE (**Supplemental Figure 2E, F**).

### Synthetic diet feeding alters the host microbiota composition and leads to systemic reductions in SCFA availability

Dietary change can drive alterations to the host intestinal microbiome, as well as metabolite availability^41,42^. Whole-genome metagenomics analysis of intestinal faecal pellets revealed significant differences between synthetic diet and chow fed mice in host microbiota composition (**Figure 3A**). Synthetic diet fed mice had an increased relative abundance of *Firmicutes* (**Figure 3B**), and significant increases in the relative abundance of *Clostridia* genera, including members of the families *Oscillospiraceae* and *Lachnospiraceae* (**Figure 3C**), which are known to metabolise dietary fibre into short-chain fatty acids (SCFAs; Refs.^43,44^).

**Figure 3.**
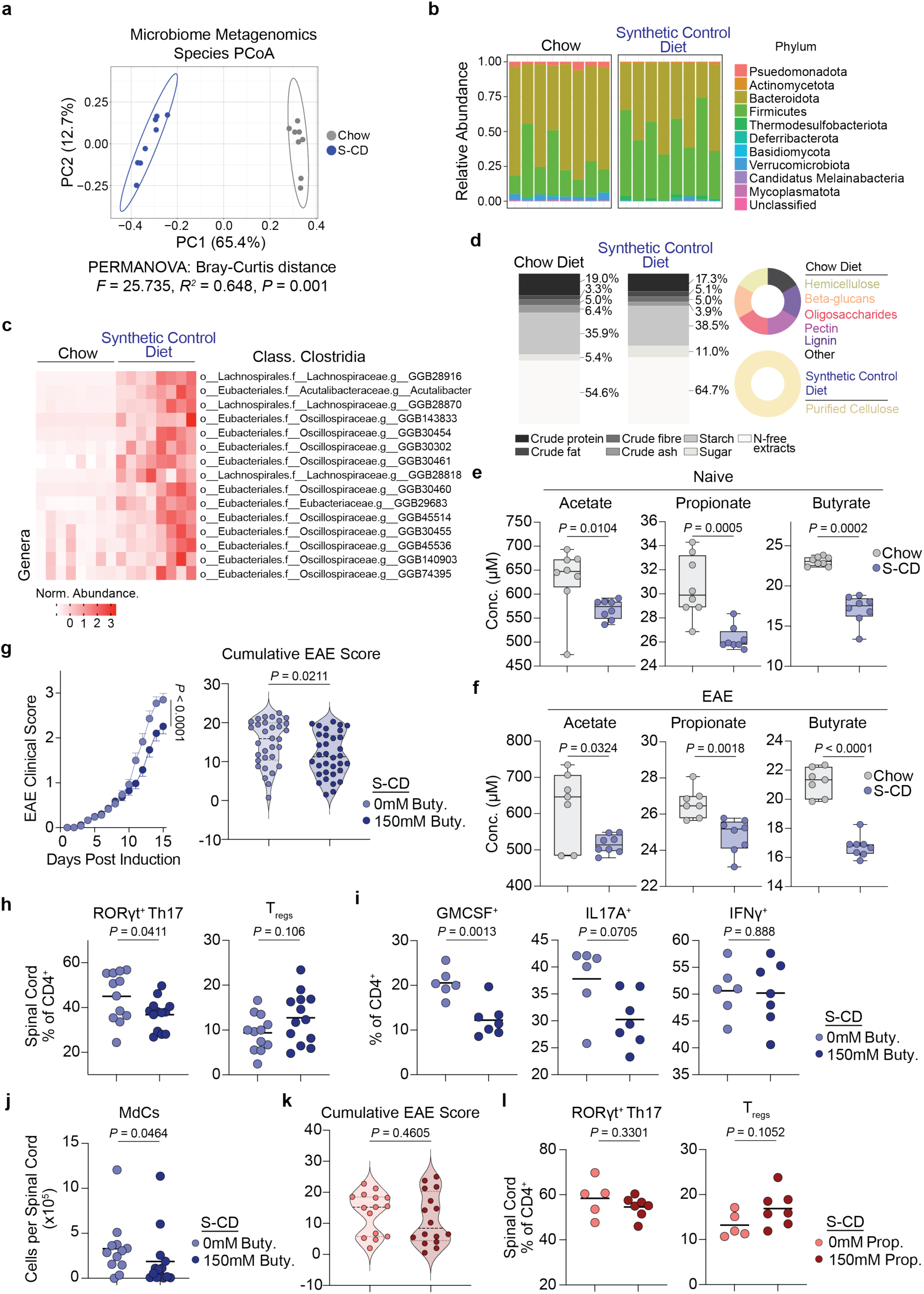
Synthetic diet feeding reprograms the host microbiome and leads to systemic SCFA deficiency. **a.** Principal coordinate analysis of microbiome diversity in small intestine faecal pellets of chow and synthetic control diet (S-CD) fed, based on species level analysis, (n = 8 per group). **b.** Relative abundance of individual bacteria phyla in the faeces of chow and S-CD fed mice. **c.** Normalised abundance of significantly differentially abundant genera from the class *Clostridia* in chow vs. S-CD fed mice. **d.** Schematic representation of nutritional composition of chow and synthetic diets. **e.** Quantification of acetate, propionate and butyrate plasma concentrations in naïve mice fed chow or S-CD. *P* values calculated using Mann Whitney U test, (n = 8 per group; EAE: n = 7 (Chow), n = 8 (S-CD)). **f.** Quantification of acetate, propionate and butyrate plasma concentrations in EAE mice fed chow or S-CD. *P* values calculated using Welch’s T-test (acetate) and parametric two-tailed unpaired T-test (propionate & butyrate). (n = 7 (Chow), n = 8 (S-CD)). **g.** Left: EAE clinical severity score over time from S-CD fed mice +/- 150mM butyrate supplementation in drinking water throughout dietary intervention. Right: cumulative EAE scores from S-CD fed mice +/- 150mM butyrate supplementation in drinking water throughout dietary intervention. Left: Error bars represent s.e.m, *P* value was calculated using two-way repeated measures ANOVA, (n = 27 per group; data from three independent experiments). Right: *P* value was calculated using Mann Whitney U Test, (n = 27 per group; data from three independent experiments). **h.** Left: relative abundance of RORγt^+^ Th17 cells among all spinal cord CD4^+^ T cells of S-CD fed mice +/- 150mM butyrate supplementation at the peak of EAE. Right: relative abundance of T_regs_ among all spinal cord CD4^+^ T cells of S-CD fed mice +/- 150mM butyrate supplementation at the peak of EAE. Black bars indicate mean values. *P* values calculated using Welch’s T-test (RORγt^+^ Th17) and parametric two-tailed unpaired T-test (T_regs_). (n = 12 (-Buty.), n = 13 (+Buty.), data from two independent experiments). **i.** Quantification of the % of GMCSF^+^ and IFNγ^+^ CD4^+^ cells from all spinal cord CD4^+^ T cells in S-CD fed mice +/- 150mM butyrate supplementation at the peak of EAE following restimulation (20nM PMA, 1µM Ionomycin, 10µg/ml Brefeldin A, 4 hours). Black bars indicate mean values. *P* values calculated using parametric two-tailed unpaired T-test (GM-CSF & IFNγ) and Mann-Whitney U test (IL-17A). (n = 5 (-Buty.), n = 6 (+Buty.)). **j.** Quantification of the number of CD45^hi^ monocyte-derived cells (MdCs) per spinal cord of S-CD fed mice +/- 150mM butyrate supplementation at the peak of EAE. Black bars indicate mean values. *P* value calculated using Mann-Whitney U test. (n = 12 (-Buty.), n = 13 (+Buty.), data from two independent experiments). **k.** Cumulative EAE scores from S-CD fed mice +/- 150mM propionate supplementation in drinking water throughout dietary intervention. *P* value calculated using parametric two-tailed unpaired T-test (n = 15 (-Prop.), n = 16 (+Prop.), data from two independent experiments). **l.** Left: relative abundance of RORγt^+^ Th17 cells among all spinal cord CD4^+^ T cells of S-CD fed mice +/- 150mM propionate supplementation at the peak of EAE. Right: relative abundance of T_regs_ among all spinal cord CD4^+^ T cells of S-CD fed mice +/- 150mM propionate supplementation at the peak of EAE. *P* value calculated using parametric two-tailed unpaired T-test (n = 5 (-Prop.), n = 6 (+Prop.)).

Unlike chow, purified cellulose, which is poorly fermented and has been shown to be poorly converted into SCFAs^45^, is the sole source of dietary fibre in synthetic diets (**Figure 3D**). Despite increases in the relative abundance of fibre metabolising microbiota, exclusive provision of dietary fibre in the form of cellulose resulted in a significant reduction in circulating concentrations of the SCFAs acetate, propionate and butyrate in synthetic diet fed mice (**Figure 3E**). These reductions were maintained during EAE, although acetate and propionate concentrations partially normalised (**Figure 3F**), while the greatest continued difference was observed in butyrate (**Figure 3F**).

### Butyrate supplementation rescues synthetic diet driven increases in EAE severity

Butyrate is a known mediator of the immune response^46^, and treatment of mice with SCFAs similar to butyrate has been shown to modulate EAE severity^47–50^. Butyrate supplementation in synthetic diet fed mice significantly reduced EAE clinical severity (**Figure 3G**). Furthermore, butyrate supplementation led to a significant reduction in the relative abundance of spinal cord RORγt^+^ Th17 cells, with a trend towards increased T_regs_ (**Figure 3H**). Butyrate supplementation further led to a significant reduction in the proportion of GM-CSF^+^ CD4^+^ T cells (**Figure 3I**), with a similar trend in IL-17A^+^ CD4^+^ T cells, and no change in the proportion of IFNγ^+^ CD4^+^ T cells (**Figure 3I**). The total abundance of spinal cord MdCs was significantly reduced following butyrate supplementation (**Figure 3J**). In contrast, propionate supplementation did not significantly affect EAE clinical severity (**Figure 3K),** and did not significantly alter the relative abundance of RORγt^+^ Th17 cells or T_regs_ (**Figure 3L**). These data suggest that systemic butyrate deficiency promotes increased EAE severity during synthetic diet feeding.

### FoxP3^+^ RORγt^+^ pT_regs_ populate the spinal cord during EAE

While the majority of T_regs_ are generated in the thymus during early life, and are referred to as “natural T_regs_” (nT_regs_), T_regs_ can also arise directly in peripheral tissues (pT_regs_) from Th cell precursors^51–59^, and can be characterised by co-expression of RORγt and FoxP3^60–62^. Critically, while butyrate has been shown to promote the expansion of intestinal pT_regs_^63–65^, whether pT_regs_ also arise in the CNS during autoimmunity and whether their abundance is affected by butyrate availability beyond the intestine remains unknown.

Profiling of 15,676 spinal cord CD4^+^ T cells from chow, S-CD and S-WD fed mice at the peak of EAE by scRNAseq revealed eight distinct clusters after removal of doublet populations and cells with high mitochondrial counts (**Figure 4A**; **Supplemental Figure 3A, B, C, D**). These clusters, present across all diets (**Supplemental Figure 3E**), included T cell subsets previously identified in the CNS of EAE mice^66^, such as migratory T cells (Clusters 0 & 2; *Cxcr6, Ccr2, Ltb4r1*), proinflammatory T cells (Cluster 1; *Ifng, Il17a, Csf2, Tnf*), as well as T_regs_ (Cluster 5; *Foxp3, Il10*) and dividing T cells (Cluster 4; *Birc5, Mki67*; **Figure 4A, B**). We also identified a subset of IFN-responsive CD4^+^ T cells (Cluster 6; **Figure 4A, B**), enriched for IFN response genes (*Ifit1, Ifit3, Isg15, Oasl2*; **Figure 4A, B**), which have previously been identified in lupus patients^67^, and in the spleen and lymph nodes of EAE mice^66^, but not spinal cord. In addition, we identified a population of homeostatic stem-like Th17 cells (Cluster 3; **Figure 4A, B**) characterised by expression of *Slamf6*, *Tcf7*, *Il6ra*, and *Ccr6* (**Figure 4A, B**), which have previously only been identified in the lymph nodes and spleen of EAE mice at disease onset, and shown to give rise to migratory pathogenic Th17 cells^66^.

**Figure 4.**
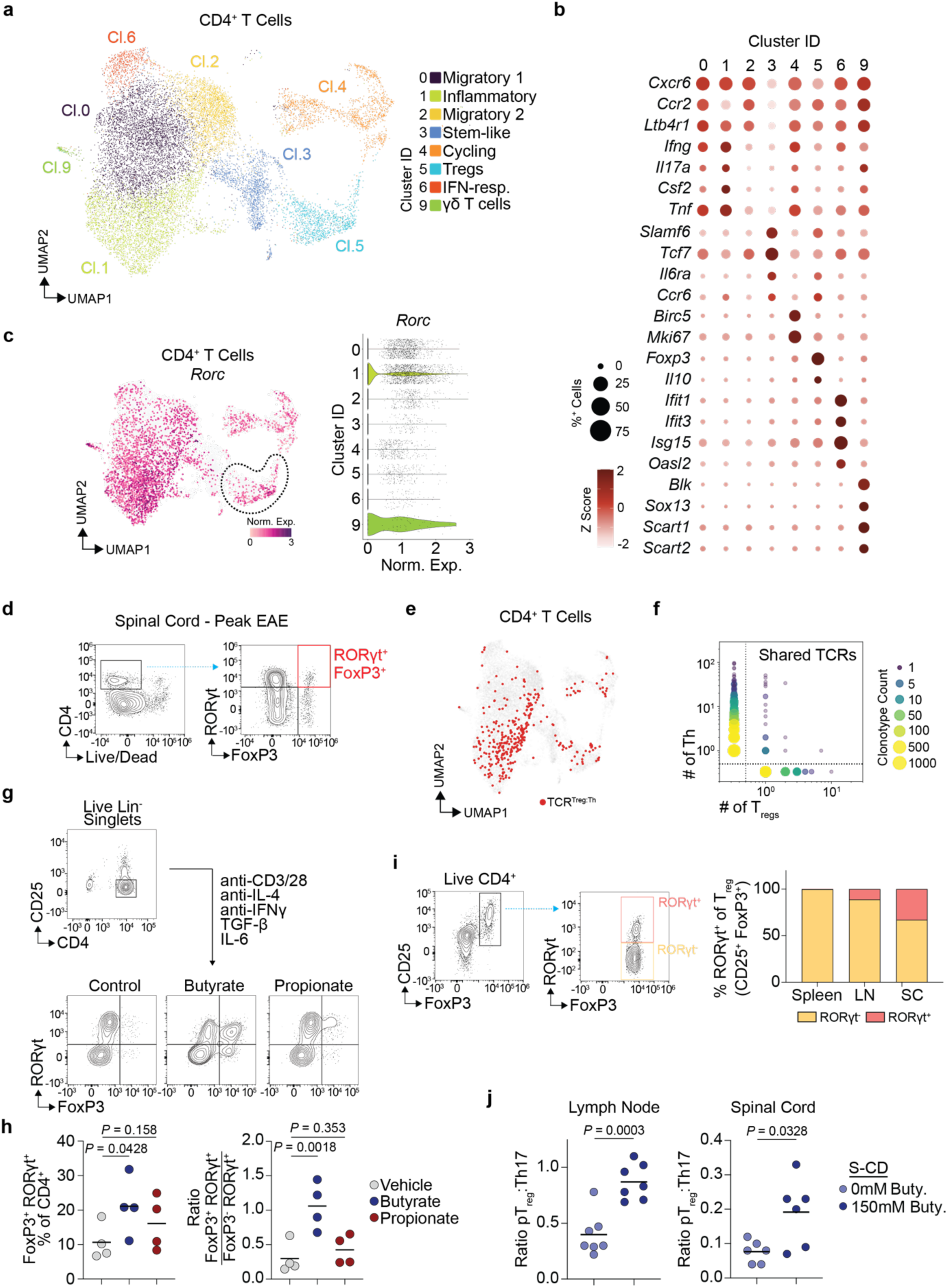
Spinal cord T_regs_ express RORγt and share TCRs with conventional Th cells. **a.** UMAP representation of unsupervised clustering of CD4^+^ T cells. **b.** Relative expression of cluster-specific marker genes. **c.** *Rorc* expression in CD4^+^ T cells. **d.** FACS based identification of RORγt^+^ FoxP3^+^ CD4^+^ T cells in spinal cord at EAE peak. **e.** UMAP representation of CD4^+^ T cells, with cells that share TCRs with T_regs_ highlighted in red. **f.** Visualisation of clonotype sharing between T_regs_ and Th cells. **g.** Gating strategy for isolation of splenic CD4^+^ CD25^-^ T cells, and gating strategy for the identification of RORγt^+^ and FoxP3^+^ cells following Th17 polarisation. **h.** Left: relative abundance of FoxP3^+^ RORγt^+^ T_regs_ among all CD4^+^ T cells following Th17 polarisation in the presence of propionate (150µM) or butyrate (150µM). Right: ratio of FoxP3^+^ RORγt^+^ T_regs_ to RORγt^+^ FoxP3^-^ Th17 cells following Th17 polarisation in the presence of propionate (150µM) or butyrate (150µM). Black bars indicate mean values. *P* values calculated using repeated-measured one-way ANOVA with Holm-Šídák’s multiple comparisons test. (n = 4 per group). **i.** Left: gating strategy for the identification of RORγt^+^ pT_regs_ and RORγt^-^ nT_regs_. Right: Proportion of RORγt^+^ and RORγt^-^ FoxP3^+^ T_regs_ from all T_regs_ (CD25^+^ FoxP3^+^) in spleen, CNS draining lymph nodes (LN), and spinal cord (SC) at EAE peak in chow fed mice (n = 6 per group). **j.** Quantification of the ratio of pT_regs_ to Th17 cells in CNS draining lymph nodes and spinal cord of S-CD fed mice +/- 150mM butyrate supplementation at the peak of EAE. Black bars indicate mean values. *P* values calculated using parametric two-tailed unpaired T-test (lymph node) and Welch’s T-test (spinal cord). (Lymph nodes: n = 7 per group; Spinal Cord, n = 6 per group).

We also identified a subset of *Rorc^+^* cells within the spinal cord T_reg_ population by scRNAseq (Cluster 5; **Figure 4C**), indicative of pT_reg_ presence, which was confirmed by FACS identification of CD4^+^ FoxP3^+^ RORγt^+^ cells in the spinal cords of mice at the peak of EAE (**Figure 4D**). Single cell TCRseq further revealed that 34 out of 831 (4.09%) TCR^+^ cells in Cluster 5 (T_regs_), which was expanded to include cycling T_regs_ previously allocated to Cluster 4 (**Supplemental Figure 3F**), shared a TCR nucleotide sequence with Th cells (**Figure 4E, F**). These data demonstrate that spinal cord T_regs_ can arise from shared clonal ancestors as T helper cells during EAE, as occurs during pT_reg_ formation.

pT_reg_ induction can be driven by RORγt^+^ antigen presenting cells (APCs; Ref. ^68–73^), as well as conventional dendritic cells (cDCs; Ref.^61,74,75^). Single cell RNAseq analysis of spinal cord myeloid cells during EAE was able to identify microglia (*Sall1^+^*, *P2ry12^+^*), monocyte-derived macrophage (*F13a1^+^*, *Tgfbi^+^*), and cDC (*Itgax^+^*, *Flt3^+^*) clusters (**Supplemental Figure 4A, B**). While the MHC class II gene *H2-Ab1* was detected in all myeloid cell clusters (**Supplemental Figure 4C**), *Rorc* was only identified in a cluster of T cell and myeloid cell doublets (**Supplemental Figure 4D**). Subclustering of cDCs revealed two predominant populations (**Supplemental Figure 4E, F**). Cluster 0 was enriched for genes associated with MHC-II antigen processing and presentation, and inflammatory response (**Supplemental Figure 4G**), while cluster 1 was enriched for migration, regulatory, and maturation genes (**Supplemental Figure 4H**). Cluster 1 cDCs therefore bore a similarity to previously characterised mature DCs enriched in immunoregulatory molecules (mregDCs), identified in tumours following antigen uptake^76^. Taken together these data show monocyte-derived macrophages are the predominant APC in the spinal cord during EAE, alongside microglia, cDC2s and mregDC-like cells. These data argue against a major contribution of spinal cord CD11b^+^*Rorc^+^* APCs to pT_reg_ induction during EAE.

### Butyrate supplementation promotes pTreg formation during EAE

Butyrate has previously been shown to promote the formation of pT_regs_ following the stimulation of naïve Th cells with anti-CD3/28 + TGF-β, through its activity as a histone deacetylase (HDAC) inhibitor^63^. Polarisation of naïve CD4^+^ CD25^-^ Th cells isolated from the spleens of naïve mice to Th17 cells, the primary pathogenic Th subset in EAE, in the presence of butyrate, led to a significant increase in the relative abundance of FoxP3^+^ RORγt^+^ pT_regs_ (**Figure 4G, H**). While propionate treatment resulted in a similar trend (**Figure 4H**), the difference was not significant, consistent with propionate being a weaker HDAC inhibitor. Butyrate treatment also significantly increased the ratio of FoxP3^+^ RORγt^+^ T_regs_ to FoxP3^-^ RORγt^+^ Th17 cells (**Figure 4H**), indicating that its ability to promote pT_reg_ formation was independent from rates of Th17 induction.

FACS analysis revealed that while pT_regs_ were largely absent from the spleen, they were robustly induced in lymph nodes and spinal cord during EAE (**Figure 4I**). Supporting our earlier *in vitro* analysis, butyrate supplementation in synthetic diet fed mice increased the relative abundance of pT_regs_ to RORγt^+^ Th17 cells *in vivo*, in both CNS draining lymph nodes and matched spinal cords during EAE (**Figure 4J**), suggesting that increased butyrate availability promotes pT_reg_ induction beyond the intestinal tract, and in the context of autoimmunity.

## Discussion

Understanding the molecular mechanisms driving diet-dependent changes to inflammation-related and autoimmune diseases is critical for identifying modifiable disease risk factors and the development of improved therapeutic interventions in an increasingly obesogenic and nutrient-deficient population. Accurate interpretation of data in murine models relies on analysing changes mediated by the specific dietary components thought to drive pathogenic responses in humans, rather than those potentially induced simply by synthetic diets alone.

By analysing the effect of synthetic diet feeding in mice, in the absence of excessive weight gain or increased dietary fats and sugars, we show that synthetic diet driven fibre deficiency reprograms the immune response during the EAE model of CNS autoimmunity, exacerbating disease severity. Mechanistically, synthetic diet feeding promotes the formation of Th17 cells, and leads to a significant increase in the expression of T-bet, which is associated with Th17 cell pathogenicity, as well as production of GM-CSF, during EAE^35–38^. Concurrently, synthetic diet fed mice display a significant increase in the relative abundance of spinal cord GM-CSF^+^ CD4^+^ T cells during EAE, and an associated increase in monocyte extravasation and differentiation into their derivate macrophages and dendritic cells. These observations reveal that synthetic diets alone are pathogenic during CNS autoimmunity.

While standard laboratory chow diets contain a diverse range of macronutrient sources, synthetic diets provide nutrition through a restricted number of purified ingredients. In particular, synthetic diet feeding and the provision of dietary fibre solely in the form of cellulose, leads to a reduction in systemic SCFA concentrations. Supplementation of synthetic diet with butyrate was able to ameliorate synthetic diet induced increases in EAE clinical severity, and correlated with a reduction in pathogenic Th17 cell induction and GM-CSF production. These data highlight the importance of using appropriate synthetic diet controls when investigating the molecular mechanisms through which dietary habits modify disease severity, and have implications for other disease models shown to be modified by dietary interventions relative to chow, such as intestinal immunity^77^ and cancer^78^, which display phenotypic rescue following SCFA supplementation.

Our data further reveals the presence of pT_regs_ in lymph nodes and spinal cord during EAE. Clonotype sharing between T helper cells and T_regs_ in EAE suggests that pT_regs_ may not be limited to expressing non-self-antigen reactive TCRs, and during EAE can instead arise from self-reactive T helper cells. Previous work has shown that pTreg induction can be driven through cDCs^61,74,75^ as well as specialist RORγt^+^ APCs^68–73^. Our analysis of the spinal cord myeloid compartment did not identify a subset of *Rorc^+^*APCs, however, not all RORγt^+^ APCs express CD11b^70^. In addition, RORγt^+^ APCs may be lowly abundant in the spinal cord, as reported in mesenteric lymph nodes^70^. Therefore, while we demonstrate that butyrate availability can augment the rate of pT_reg_ induction during EAE, future studies should aim to mechanistically resolve whether CNS pT_reg_ specification is driven by CNS-resident or peripherally-derived APCs.

Finally, the increased consumption of ultra-processed foods (UPFs), which similar to murine synthetic diets are nutrient deficient, particularly in fibres^79,80^, is an emerging factor in the development of multiple diseases such as cancer, type-II diabetes, cardiovascular disease and Chron’s disease^81^. Due to the recent emergence of UPFs, studies remain limited in their scope, however elements of UPFs have been linked to chronic inflammation^82^, and UPF driven reductions in SCFA abundance have been posited to exacerbate UPF driven immune disturbances and promote inflammation^83^. Our data opens the possibility that the consumption of UPFs, and subsequent reduction in SCFA abundance, may dysregulate the immune response in humans, potentially through altering the induction of pathogenic T cell subsets as well as anti-inflammatory pT_regs_.

## Limitations of the study

While we are able to show that synthetic diet exposure reduces the relative abundance of spinal cord T_regs_, we do not know whether the remaining T_regs_ following synthetic diet feeding are qualitatively different to their chow counterparts. Differences in T_reg_ signaling could contribute to diet-specific alterations in Th17 cell abundance and EAE pathology. In addition, analysis of the myeloid cell compartment revealed multiple APC populations, although no evidence of a specific *Rorc^+^* population. This could be due to *Rorc^+^* APCs being absent from the spinal cord or negative for CD11b expression, and would require a more comprehensive scRNAseq with a larger cell number input.

**Supplemental Figure 1.**
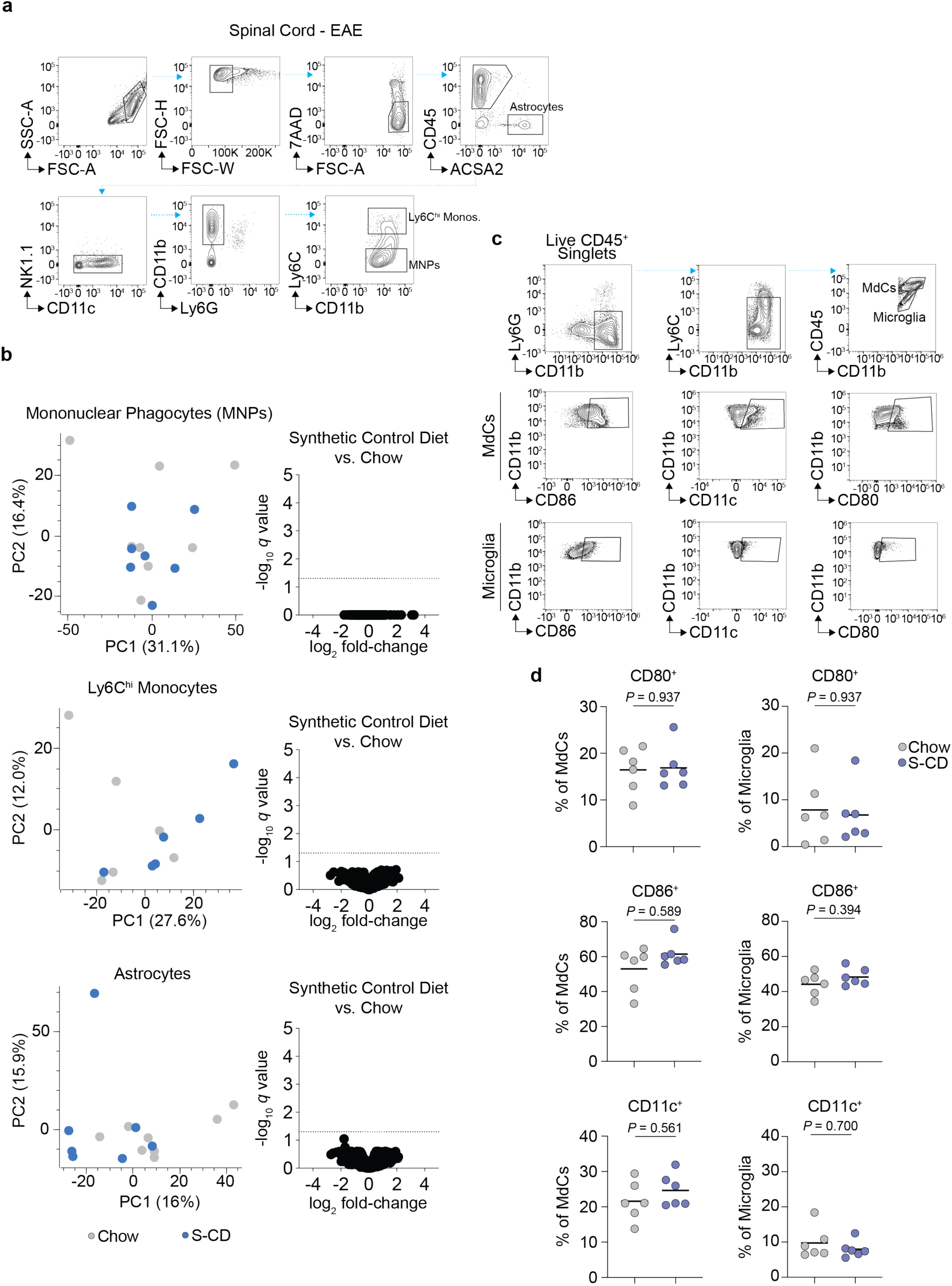
LC-MS/MS analysis of spinal cord myeloid cells and astrocytes from synthetic diet and chow fed mice. **a.** Gating strategy for isolation of astrocytes, Ly6C^hi^ monocytes and mononuclear phagocytes (MNPs). **b.** Volcano plots of protein expression fold-change versus Adj. *P*-values (*q*-value) in LC-MS/MS analysis of mononuclear phagocytes (MNPs), Ly6C^hi^ monocytes and astrocytes at EAE peak from mice fed S-CD or chow. Dotted line indicates *q* = 0.05. **c.** Gating strategy for analysis of spinal cord monocyte-derived cells (MdCs) and microglia. **d.** Quantification of the relative abundance of CD80^+^, CD86^+^ and CD11c^+^ MdCs and microglia in S-CD fed mice relative to chow at EAE peak. Black bars indicate mean values, *P* values calculated using Mann Whitney U test.

**Supplemental Figure 2.**
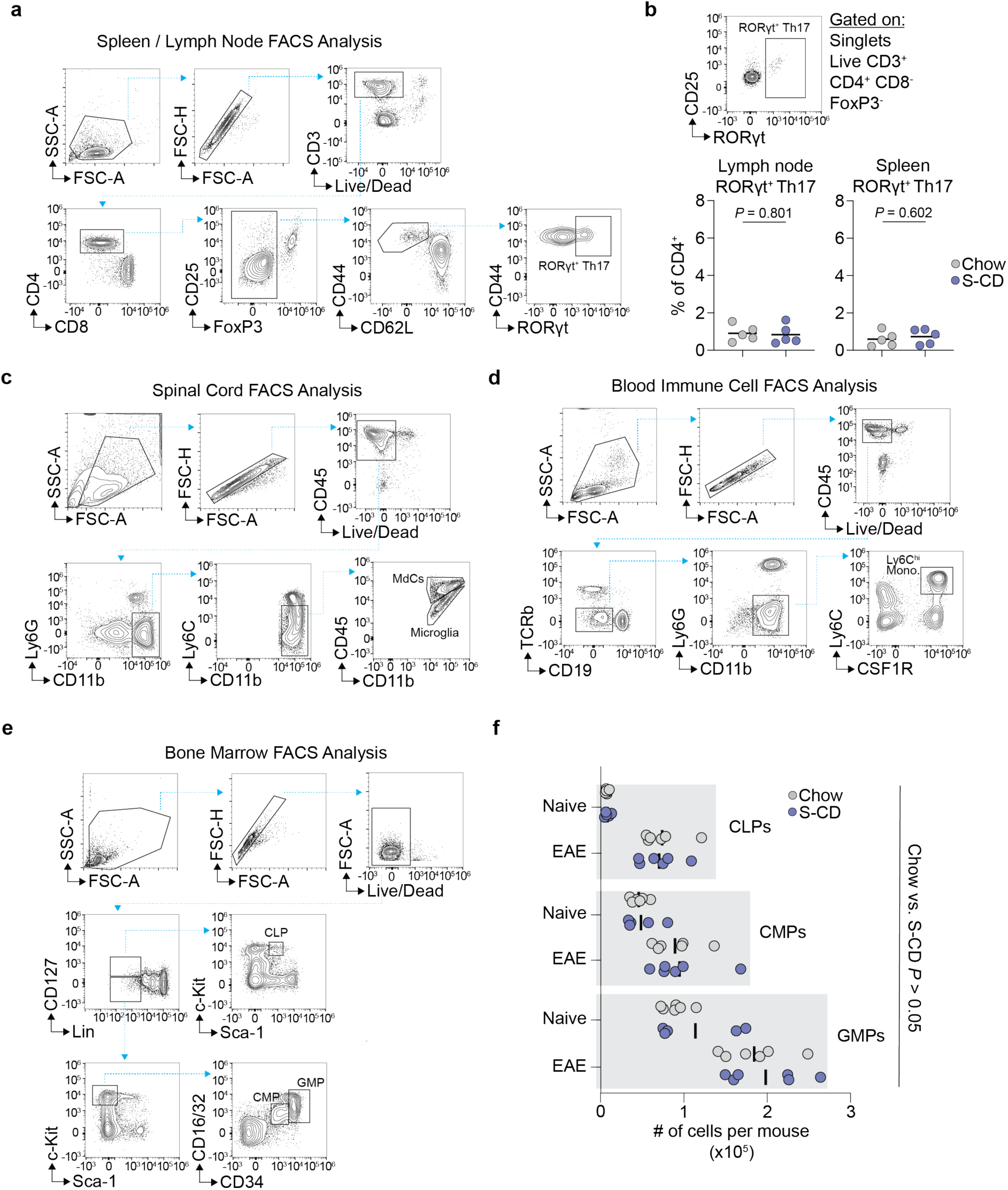
FACS gating strategies for the analysis of lymph node, spleen, spinal cord, blood and bone marrow cell abundances. **a.** Gating strategy for the analysis of RORγt^+^ Th17 cells in lymph nodes and spleen. **b.** Quantification of the relative abundance of RORγt^+^ Th17 cells in inguinal lymph nodes and spleen from mice fed S-CD or chow 5 days after immunisation with complete freund’s adjuvant in the absence of MOG_35-55_. **c.** Gating strategy for the analysis of Ly6C^hi^ monocytes, monocyte-derived cells (MdCs) and microglia in spinal cords. **d.** Gating strategy for the analysis of circulating Ly6C^hi^ monocytes in blood. **e.** Gating strategy for the identification of immune progenitor subsets in bone marrow (GMP = granulocyte-monocyte progenitor, CLP = common lymphoid progenitor, CMP = common myeloid progenitor). **f.** Quantification of the abundance of immune progenitor subsets in S-CD and chow fed naïve mice and during EAE onset. All comparisons *P* > 0.05 as determined by Mann Whitney U test (n = 6 to 11).

**Supplemental Figure 3.**
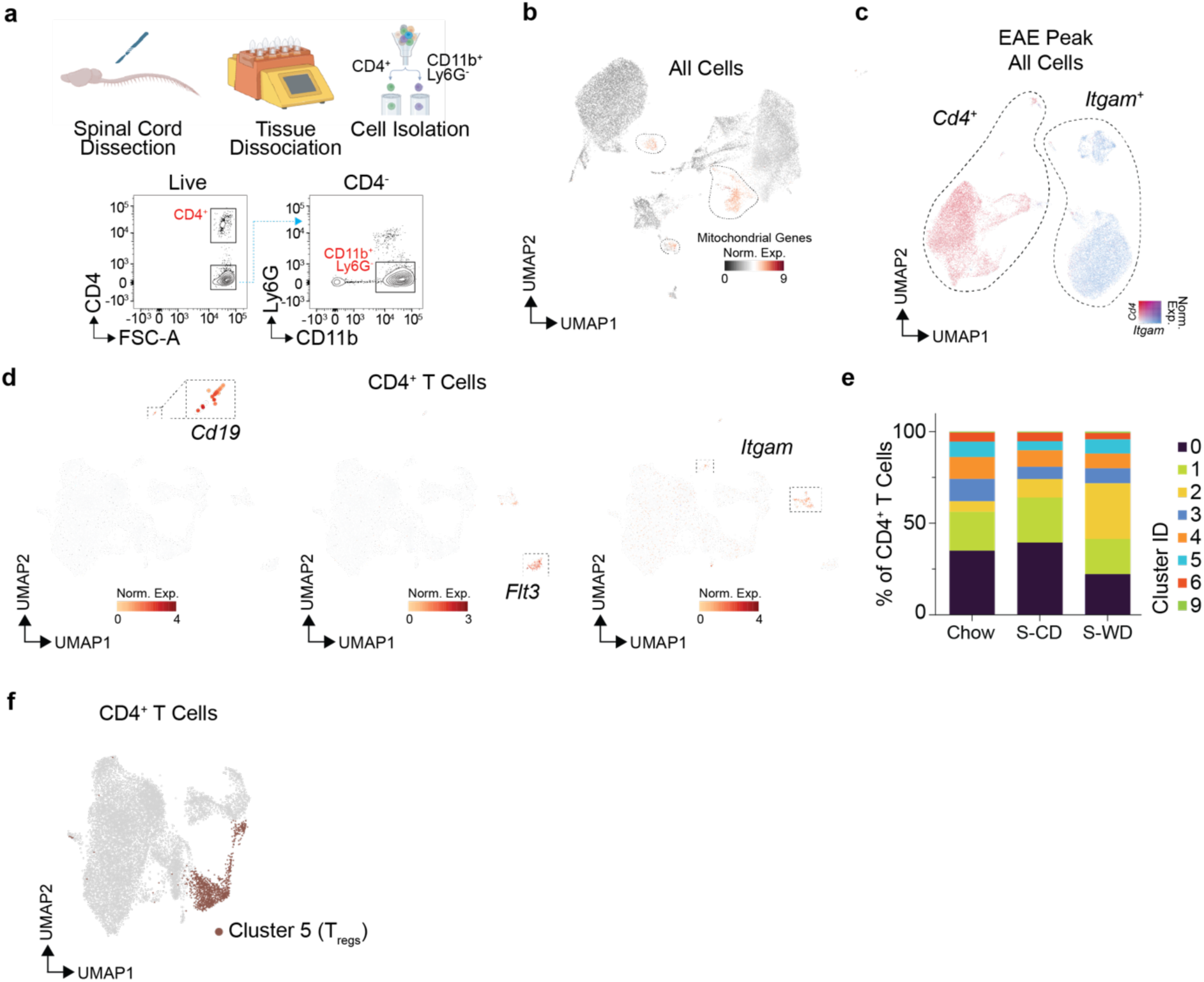
Spinal cord CD4^+^ T cell scRNAseq. **a.** Schematic representation of CD4^+^ T cell and CD11b^+^ Ly6G^-^ myeloid cell isolation for scRNAseq. Created in BioRender. Weiss, F. (2026). **b.** UMAP representation of CD4^+^ and CD11b^+^ Ly6G^-^ cells following scRNAseq, with mitochondrial gene expression highlighted in red. Selected cell clusters were removed for downstream analysis. **c.** Identification of CD4^+^ T cells for downstream analysis **d.** Identification of doublets in CD4^+^ T cell subset based on *Cd19*, *Flt3* and *Itgam* expression. Selected cell clusters were removed for downstream analysis. **e.** Relative abundance of each T cell cluster in chow (n = 4), S-CD (n = 2) and S-WD (n = 2) fed mice. **f.** Clustering of T_regs_ used for clonotype sharing analysis to including cycling T_regs_.

**Supplementary Figure 4.**
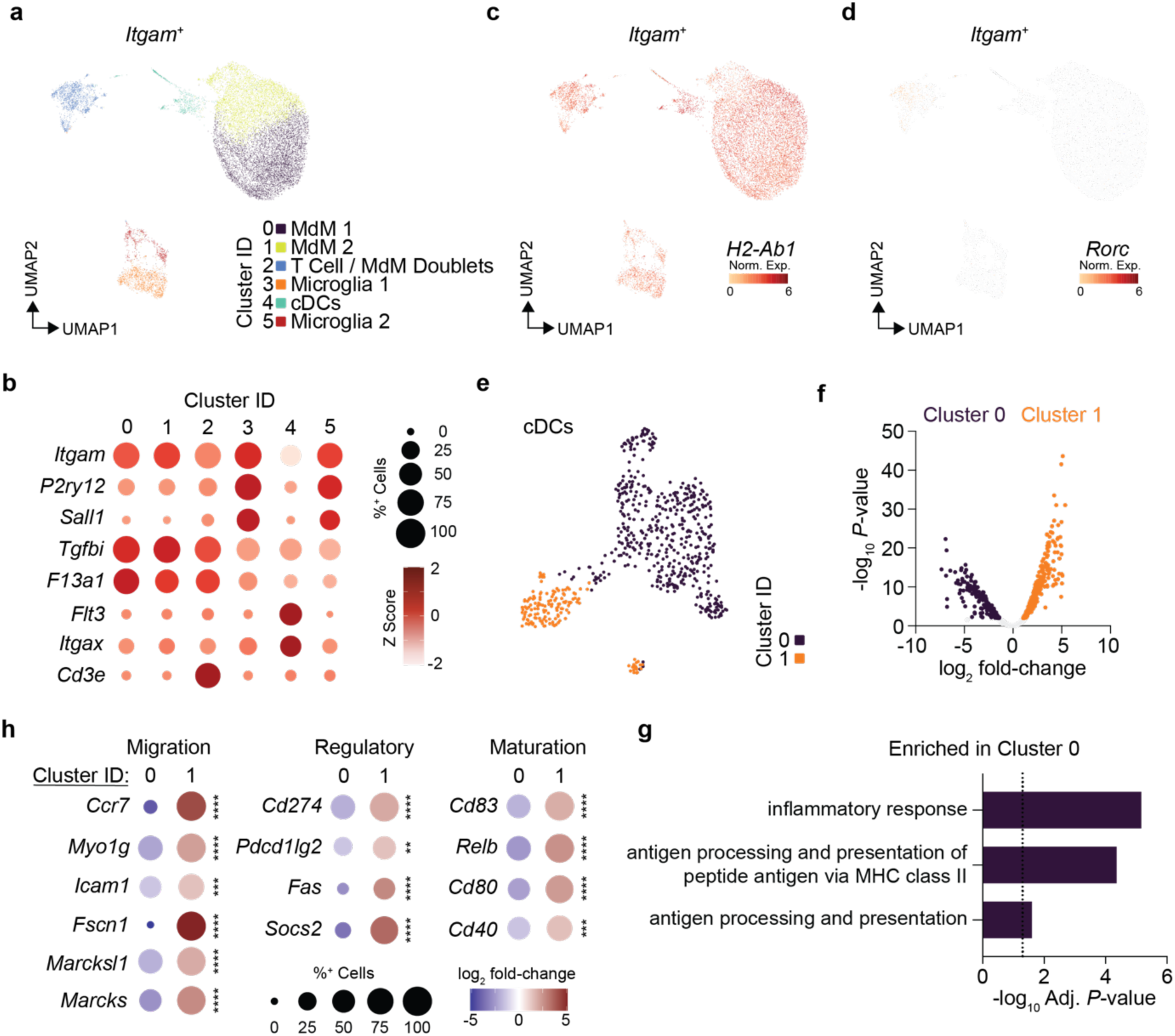
Spinal cord myeloid cell scRNAseq. **a.** UMAP representation of unsupervised clustering of CD11b^+^ Ly6G^-^ myeloid cells. **b.** Relative expression of cluster-specific marker genes. **c.** *H2-Ab1* expression in myeloid cells. **d.** *Rorc* expression in myeloid cells. **e.** UMAP representation of unsupervised clustering of conventional dendritic cells (cDCs, Myeloid Cluster 4). **f.** *P* value versus fold-change of gene expression differences between cDC cluster 0 and cluster 1. Genes with FDR > 0.01 shown in grey, genes significantly enriched (FDR < 0.01, log_2_ fold-change < 0) in cluster 0 shown in blue, genes significantly enriched (FDR < 0.01, log_2_ fold-change > 0) in cluster 1 shown in orange. *P* values calculated using negative binomial test, and adjusted for multiple testing using Benjamini-Hochberg. **g.** Significantly enriched gene ontology (biological process) terms for genes significantly enriched in cDC cluster 0 relative to cluster 1. Adjusted *P* value calculated using g:SCS algorithm from gProfiler. **h.** Relative expression of migration, regulatory and maturation associated genes in cDC clusters 1 and 2. ** FDR < 0.01, *** FDR < 0.001, **** FDR < 0.0001, *P* values calculated using negative binomial test, and adjusted for multiple testing using Benjamini-Hochberg.

## Materials & Methods

### Animal Housing

C57B6/J mice were obtained from Charles River Laboratories at six weeks of age, and housed for a minimum of two weeks before the start of experimental interventions. Mice were maintained under specific pathogen-free (SPF) conditions at the Immunological Facility of Experimental Therapy (iFET, University Hospital Bonn). Animals were given water ad libitum and kept under a standard 12-hour circadian light/dark cycle. All experiments were performed in accordance with the German law of animal protection and the Landesamt für Natur, Umwelt und Verbraucherschutz (LANUV) and with permission of the state of Nordrhein-Westfalen.

### EAE Induction and Scoring

For EAE induction, male C57B6/J mice were subcutaneously injected at the tail base with 50µg MOG_35-55_ peptide (Genscript, RP10245) and 0.2mg heat-killed *Mycobacterium tuberculosis* (BD Biosciences, 231141) emulsified in complete Freund’s adjuvant (CFA; Sigma Aldrich, F5881). 200 – 400ng Pertussis toxin (Merck, 516560) was injected intraperitoneally on day 0 and day 2 after initial immunisation. Clinical signs of EAE were assessed along a scale as follows: 0, no clinical signs; 1, complete loss of tail tonus; 2, partial hindlimb paresis; 3, bilateral hindlimb paresis; 4, complete fore- and hindlimb paresis. Mice which failed to develop EAE (EAE score < 0.50) were excluded from analyses and experiments. For CFA only controls, mice were immunised as described above, in the absence of MOG_35-55_ peptide. For dietary intervention studies, 8-week-old mice were either maintained on chow (Ssniff, V1534-300) or swapped to γ-irradiated synthetic control diet (Ssniff, CD.88137) or synthetic Western diet (Ssniff, TD.88137), and fed for a period of 8 weeks prior to EAE induction, and then maintained on the same diets throughout EAE. For SCFA supplementation, mice were provided drinking water supplemented with 150mM Sodium Butyrate (Sigma, 303410) as previously described^84^, or 150mM Sodium Propionate (Sigma, P1880), throughout synthetic diet feeding and during EAE. SCFA supplemented water was replaced fresh every 48 to 72 hours. For all analyses, EAE onset was defined as 7 days post-immunisation and EAE peak as 14-16 days post-immunisation. Cages of co-housed males were randomly assigned to experimental conditions.

### FACS Analysis and Isolation of Immune Cells

Mice were sacrificed by intraperitoneal administration of Ketamine (100mg/kg) / Xylazine (20mg/kg), and blood drawn from the vena cava prior to transcardiac perfusion with ice cold PBS. Blood was collected with 10µl of 0.5M EDTA, and subject to two rounds of red blood cell lysis (Thermofisher, 00-4333-57), before antibody staining. Following perfusion, bone marrow from the tibia and femur was flushed with PBS + 2mM EDTA, and the lumbar aortic and inguinal lymph nodes and spleen were excised, and minced through an 85µm mesh to obtain a single cell solution. Spleens were subjected to two rounds of red blood cell lysis (Thermofisher, 00-4333-57), before antibody staining. Subsequently, whole spinal cord was removed and subjected to digestion and myelin removal according to manufacturer’s instructions (Neural Tissue Dissociation Kit (P), Miltenyi Biotec, 130-092-628; Myelin Removal Kit II, Miltenyi Biotec, 130-096-433).

For the isolation of cells from the CNS, single cell suspensions were stained with: ACSA-2 PE (Miltenyi Biotec, 130-123-284), CD45 PE-Cy7 (BD Biosciences, 552848), NK1.1 APC (Biolegend, 108709), CD11c BV510 (Biolegend, 117337), CD11b FITC (Biolegend, 101205), Ly6G BV421 (Biolegend, 127628), Ly6C APC-Cy7 (Biolegend, 128025). For the FACS analysis of spinal cord myeloid cells, single cell suspensions were stained with: CD45 PE-Cy7 (BD Biosciences, 552848), Ly6G PE/Dazzle 594 (Biolegend, 127647), CD11b FITC (Biolegend, 101205), Ly6C PerCP-Cy5.5 (Biolegend, 128027).

For the FACS analysis of bone marrow immune progenitor cells, single cell suspensions were stained with: BD Horizon lineage cocktail V450 (BD Biosciences, 561301), CD127 BV650 (Thermofisher, 25-1271-82), c-Kit FITC (BD Biosciences, 561680), Sca-1 BV786 (BD Biosciences, 563991), CD34 PE-Cy7 (Biolegend, 128618), CD16/32 BV711 (Biolegend, 101337).

For the FACS analysis of blood cells, single cell suspensions were stained with: CD45 PE-Cy7 (BD Biosciences, 552848), TCRb BUV615 (BD Biosciences, 751212), CD19 FITC (Biolegend, 115506), Ly6G BV421 (Biolegend, 127628), CD11b BV510 (Biolegend, 101245), Ly6C PerCP-Cy5.5 (Biolegend, 128027), CD115 PE (Biolegend, 135506).

For the FACS analysis of T cell transcription factors from all tissues, cells were initially stained with CD3 APC (Biolegend, 100236), CD4 PerCP Cy5.5 (Biolegend, 100433), CD8 BV570 (Biolegend, 100739), CD25 BV421 (Biolegend, 102033), CD44 BV785 (Biolegend, 103041), CD62L BV605 (Biolegend, 104438), and then fixed and permeabilised according to manufacturer’s instructions (FoxP3 / Transcription Factor Staining Buffer Set, ThermoFisher Scientific 00-5523-00), and subsequently stained with FoxP3 AF488 (Biolegend, 320012) and RORγt RB744 (BD Biosciences, 570490). T-bet PE/Fire 810 (Biolegend, 644839) was stained overnight at 4°C. For the quantification of pT_reg_ to Th17 ratios, samples with fewer than 50 total detected T_regs_ were excluded.

For the FACS analysis of cytokines, CD4^+^ T cells were enriched from single cell suspensions according to manufacturer’s instructions (CD4^+^ T Cell Isolation Kit, Miltenyi Biotec 130-104-454) and stimulated in 20nM PMA (Sigma, P8139), 1µM Ionomycin (Invitrogen, I24222)) and 10µg/ml Brefeldin A (Biolegend, 420601) for 4 hours, prior to antibody staining. For the FACS analysis of T cell cytokines, cells were initially stained with CD90.2 PE/Dazzle 594 (Biolegend, 140328), CD4 PerCP Cy5.5 (Biolegend, 100433), CD8 BV570 (Biolegend, 100739), CD44 BV785 (Biolegend, 103041), CD62L BV605 (Biolegend, 104438), and then fixed and permeabilised according to manufacturer’s instruction (FoxP3 / Transcription Factor Staining Buffer Set, ThermoFisher Scientific 00-5523-00), and subsequently stained with IL-17A BV650 (Biolegend, 506930), GMCSF FITC (Biolegend, 505404), IFNγ BV510 (Biolegend, 103043).

With the exception of bone marrow, all samples were co-incubated with anti-mouse CD16/32 Fc Block (BD Biosciences, 553141) during antibody staining. Live/dead discrimination was performed with 7AAD (BioLegend, 420404) or Fixable Near-IR stain (ThermoFisher, L34975) according to manufacturer’s instructions. Stained cells were resuspended in FACS buffer, and counting beads (Biolegend, 424902) added where appropriate. Cell numbers were calculated according to manufacturer’s instructions. Cells were analysed on Sony ID7000 5L, and sorted with the BD Aria III or BD Fusion.

### T Cell Polarization

The day before T cell isolation, a flat bottom 96-well plate was coated with 2µg/ml α-CD3 (Biolegend, 100340) and 2µg/ml α-CD28 (Biolegend, 102116) diluted in PBS and incubated overnight at 4°C. Inguinal lymph nodes and spleen were harvested and a single cell suspension was generated as described previously. CD4^+^ T cell enrichment was performed following the manufacturer’s instructions (Miltenyi, 130-096-533). 4x10^4^ cells were seeded in RPMI (Gibco, 61870044), supplemented with 10% FCS, 10mM HEPES (Gibco, 15630080), 100U/ml Pen/Strep (Merck, P4333), 1mM Sodium Pyruvate (Gibco, 11360-070), 1x NEAA (Gibco, 11140050), 50µM 2-Mercaptoethanol (ThermoFisher, 35602) and polarized for 3.5 days at 37°C and 5% CO_2_ using 10µg/ml α-IL-4 (Biolegend, 504135), 10µg/ml α-IFNγ (Biolegend, 505847), 50ng/ml mIL-6 (Biolegend, 2116693) and 5ng/ml hTGF-ß1 (Biolegend, 781802) in the presence or absence of 150µM Sodium Butyrate (Sigma, 303410) or 150µM Sodium Propionate (Sigma, 303410).

### Immunofluorescence Analysis

30 days following EAE induction mice were sacrificed by intraperitoneal administration of Ketamine (100mg/kg) / Xylazine (20mg/kg), and perfused with ice cold PBS followed by 4% paraformaldehyde (PFA, ThermoFisher, J19943.K2). Extracted spinal cords were post-fixed in 4% PFA at 4°C overnight, rinsed in PBS and transferred in PBS + 30% glucose solution at 4°C. Samples were subsequently embedded in Tissue-Tek O.C.T. (Diagonal, 1115200), using cryomolds (VWR, IBSA4566) and stored at -80°C. 12µm lumbar, thoracic and cervical sections were sliced on to polylysine slides (ThermoFisher, 50-279-88). Samples were washed with TBS prior to blocking and permeabilisation with blocking solution (0.5% Triton-X-100 and 10% FCS in TBS) for 1 hour. MBP-antibody (Bio RAD, MCA409S) solution was prepared in blocking solution and incubated overnight at 4°C. Subsequently, secondary anti-rat Alexa Fluor 647 (Invitrogen, A-212-7) antibody staining was performed for one hour at room temperature. Stained samples were washed and mounted with DAPI-containing Fluoromount-G Mounting Medium (ThermoFisher, 00-4958-02). Samples were imaged on a Zeiss AxioscanZ.1, with Zeiss ZEN (v3.5). Lesions were defined as spinal cord regions enriched for DAPI and absent for MBP.

### Metagenomics Library Preparation and Sequencing

Faecal pellets were removed from the small intestine and DNA extracted using PureLink Microbiome DNA Purification Kit (Invitrogen, A29790) according to manufacturer’s instructions. DNA samples were quantified using Qubit 4.0 Fluorometer (Life Technologies, Carlsbad, CA, USA) and DNA integrity was checked using Agilent TapeStation 4200 (Agilent Technologies, Palo Alto, CA, USA). NEBNext Ultra II DNA Library Preparation kit for was used for DNA library preparation following the manufacturer’s recommendations (NEB, Ipswich, MA, USA). Briefly, genomic DNA was fragmented by acoustic shearing with a Covaris instrument. Fragmented DNA was end repaired and adenylated. Adapters were ligated after adenylation of the 3’ends followed by enrichment by limited cycle PCR. Adapter-ligated DNA libraries were cleaned up and validated using Agilent TapeStation and quantified using Qubit 4.0 Fluorometer. The libraries have been multiplexed on a flowcell and loaded on the Illumina NovaSeq X plus instrument according to the manufacturer’s instructions. The samples were sequenced using a 2x150 paired-end (PE) configuration. Image analysis and base calling was conducted by the NovaSeq Control Software v1.3.1 on the NovaSeq instrument. Raw sequence data (.bcl files) generated from Illumina NovaSeq was converted into fastq files and de-multiplexed using Illumina bcl2fastq program version 2.20. One mismatch was allowed for index sequence identification.

### Metagenomics Analysis

Preprocessing and quality control of the sequenced samples were performed as before^85^. Taxonomic profiling was performed using MetaPhlAn (version 4.1.0)^86^. Beta diversity analyses were performed in R (v4) using Bray–Curtis dissimilarities calculated from relative abundance tables at the phylum and species levels. Differences in microbial community composition across diet groups were assessed using permutational multivariate analysis of variance (PERMANOVA) implemented in the adonis2 function from the vegan package with 999 permutations. Homogeneity of multivariate dispersion was evaluated using the betadisper function followed by permutation testing (999 permutations) to ensure that significant PERMANOVA results were not driven by differences in within-group dispersion. Pairwise PERMANOVA comparisons between diet groups were conducted with Benjamini–Hochberg correction to account for multiple testing.

### Short-chain Fatty Acid Quantification

A single-phase extraction protocol using 2-propanol (IPA) was applied for metabolite extraction from stool and plasma samples. Stool samples were first homogenized in IPA to ensure thorough disruption and solubilization of metabolites. For each extraction, 50μL of stool or plasma was mixed with 100μL of an internal standard solution containing 13C, D3-acetate, D5-propionate, and D7-butyrate (10μg/mL) was added. IPA was then introduced to achieve a final concentration of 70%. The samples were incubated at 4°C for 60 minutes, followed by centrifugation at 3500 × g for 10 minutes at 4°C. The supernatant, containing the metabolites, was transferred to a new glass vial^87^. Derivatization was performed using 3-nitrophenylhydrazine hydrochloride and N-(3-dimethylaminopropyl)-N′-ethylcarbodiimide hydrochloride (Sigma–Aldrich, Saint Louis, MO, USA), as previously described^88^. A 10µL aliquot of the derivatized sample was then transferred to a 384-well plate. The samples were analysed by direct infusion mass spectrometry (DI-MS) using a Triversa Nanomate (Advion BioSciences Inc.) source coupled with a Q Exactive-HF Hybrid Quadrupole-Orbitrap mass spectrometer (Thermo Scientific, Bremen, Germany)^89^.

The TriVersa NanoMate^TM^ ion source was run in negative modus with the voltage of 1.2 kV and the head pressure of 0.25 psi. 5µl of sample were injected into the mass spectrometer for 2 min. The mass spectrometer was operated in negative modus with the capillary temperature set to 320°C and the RF level to 55 %. The instrument was set to acquire both Full MS1 and MS2 (PRM) spectra of the selected precursors. The MS1 scan was done in a range from 50 to 300 m/z with an orbitrap resolution of 120,000 at 200 m/z (AGC 3e6, maximum injection time 200ms). The targeted PRM scans were carried out with the following settings: orbitrap resolution 30,000, maximum injection time 100ms, AGC 2e5, isolation window 1.0 m/z, normalised collision energy (NCE) 30%. The DI-MS data were manually curated using FreeStyle software (Thermo Fisher Scientific, Rockford, USA).

### Single cell RNAseq

Single-cell RNA sequencing (scRNA-seq) libraries were generated using the 10x Genomics GEM-X Universal 5′ Gene Expression v3 4-plex and Chromium Single Cell Human TCR Amplification Kit with on-chip multiplexing (OCM), following the manufacturer’s instructions (CG000770 Rev. A; 10x Genomics). The GEM-X Universal 5′ Gene Expression v3 4-plex assay provides a scalable microfluidic platform for OCM of up to eight samples (two sets of four samples each), enabling simultaneous assessment of gene expression and immune repertoire from the same single cells. Briefly, single-cell suspensions of spinal cord isolated CD4^+^ and CD11b^+^ Ly6G^-^ cells mixed at a 1:1 ratio were prepared and adjusted to a concentration of 600–1300 cells per µl. A total of 5,000 cells per sample were loaded onto a GEM-X OCM chip for multiplexing and Gel Beads-in-emulsion (GEM) generation. Barcoded cDNA was purified and amplified, followed by V(D)J enrichment and separate library construction for gene expression (GEX) and V(D)J profiling. Dual-indexed libraries were quantified, pooled, and sequenced on an Illumina NovaSeq X Plus platform, targeting a minimum sequencing depth of 20,000 reads per cell for GEX libraries and 10,000 reads per cell for V(D)J libraries.

### Single cell RNAseq Analysis

Raw sequencing reads were processed with cellranger-count (v9.0.1). Demultiplexing, read mapping, detection of intact cells, and gene expression quantification were performed using the cellranger –multi pipeline, including both the gene expression and V(D)J libraries, with default parameters and refdata-cellranger-arc-GRCm39-2024-A and refdata-cellranger-vdj_GRCm38_alts_ensembl-mouse-2.2.0 as reference. This yielded to 3597, 5218, 2485, 4900, 3413, 5753, 4815, 4326 for Chow_1, SCD_1, Chow_2, Chow_3, Chow_4, SWD_1, SWD_2, SCD_2, respectively. The libraries were then merged with cellranger-aggr without library normalization. Further analysis was performed using R (v4.4.1) and Seurat (v5.1.0). Briefly, the gene-expression matrix was loaded with Read10X_h5, immunoglobulin transcripts were removed (based on the biotype annotation in the reference gene.gtf file), and a Seurat object was created with CreateSeuratObject. A Uniform Manifold Approximation and Projection (UMAP) was computed using ScaleData, RunPCA (50 PCs), and RunUMAP with dimensions 1:50. Low-quality clusters were identified through visual inspection of mitochondrial gene content, total UMI counts as well as the number of detected genes and removed from further analysis (**Supplemental Figure 3B**). T cells were separated from myeloid cells by re-embedding the remaining cells in a new UMAP and projecting the expression of CD4 and ITGAM (**Supplemental Figure 3C**). Myeloid, cDC and CD4^+^ T cells were then re-analysed independently to generate a dedicated UMAPs. Clusters with similar transcriptional profile were identified using shared-nearest-neighbor (SNN) modularity optimisation via FindNeighbors (PCA reduction, dimensions 1:50) and FindClusters with resolutions ranging from 0.1 to 1.0 in 0.1-step increments. For T cell data, a resolution of 0.3 was chosen as optimal based on the expression of selected marker genes. Clusters 7, 8, and 10, exhibiting expression of *Cd19*, *Flt3*, and *Itgam*, were likely doublets or contaminants and excluded from downstream analysis (**Supplemental Figure 3D**). For myeloid cell analysis a resolution of 0.2, and for dendritic cells a resolution of 0.1, was chosen as optimal based on the expression of selected marker genes. Differential expression was performed with Loupe Browser (v9.1, 10X Genomics), and only genes with a mean expression of 1 read per cell were included. Gene ontology analysis was performed with gProfiler, with a custom list of all expressed genes in each comparison used as background.

### Single cell TCRseq Analysis

For the TCR repertoire analysis, clonotypes were defined using the full nucleotide V(D)J sequence of the alpha and beta chains provided by cellranger (v9.0.1). For each locus a primary and secondary chain, when present, were annotated onto cells with scirpy (v0.12.1). Cells with incomplete nucleotide sequences were removed (27), specifically due to missing CDR2 annotation for V genes TRAV9D-4 or TRBV21. While maintaining nucleotide resolution for clonotype definition, for practicality and plotting each sequence was summarized with its corresponding bioidentity, defined as the CDR3 amino acid sequence followed by the V gene name. Inconsistent V gene annotations for the same sequence were corrected to a unique bioidentity using IMGT HighV-Quest (164 TRB clonotypes from TRBV12-2 or TRBV13-1 reannotated as TRBV13-2, and 3 TRA clonotypes from the families TRAV14N and TRAV14D reannotated as TRAV14-2). However, 66 bioidentities corresponding to multiple nucleotide sequences (2-8) were retained since sequencing errors or convergence at amino acid level are expected to occur. Finally, to remove potential artifacts, clonotypes were filtered allowing at most one secondary chain, either from TRA or TRB. For clonotype sharing analysis, T_reg_ Cluster 5 was expanded to include cycling T_regs_, and Th cells were defined as cells in all clusters except Cluster 5.

### LC-MS/MS Sample Preparation

Snap-frozen sorted cell pellets were denatured in Radioimmunoprecipitation (RIPA, NaCl 150mM, Tris-HCl 50mM pH 7.5, 1% SDC, 0.1% SDS, 1% Triton X-100) buffer and proteins were reduced and alkylated with 10mM tris(2-carboxyethyl)phosphine (BondBreaker^TM^, Thermofisher, B3026-77720) and 30mM 2-Chloroacetamide in the dark for 10 min at 90°C while shaking. Proteins were purified and digested by Protein Aggregation Capture (PAC, Ref. ^90^) on a Bravo automated liquid handling platform. Per sample, 1.5µl MagReSyn® Hydroxyl magnetic microparticles (20mg/ml suspension) were washed twice in 70% acetonitrile (ACN; LC-MS Chromasolv^TM^) with help of a magnet before adding 50µl lysate. ACN was added to a final concentration of 70% and samples were incubated at RT for 10 min to allow proteins to aggregate on the magnetic nanoparticles. Beads were sequentially washed with 100% ACN and 70% ethanol and proteins were digested with 100mM Tris/HCl buffer (pH 8.5) containing 150ng (5ng/µl) Trypsin and LysC per sample at 37°C for 5 hours, 800 rpm. Digestion was quenched with a final of 0.1% trifluoracetic acid (TFA). Peptides were collected and combined with a second wash of the beads using 0.1% TFA. Peptides were loaded and de-salted on C18 solid phase disposable trap columns (Evotip One columns, Evosep Biosystems), following the manufacturer’s protocol.

### LC-MS/MS Measurements

Astrocytes, Ly6C^hi^ monocytes, and MNPs were measured independently, and sample measurements were randomised to avoid confounding effects. Peptides were separated by reverse-phase liquid chromatography (LC) with an Evosep One on a 15cm C18 UHPLC column (Aurora Elite CSI, Ionopticks) using a binary buffer system of 0.1% formic acid (FA) (Buffer A) and ACN, 0.1% FA (Buffer B) using the pre-installed Whisper 40 samples-per-day non-linear gradient at a constant flow rate of 100nl/min. Column temperature was monitored and maintained at 50°C. Eluting peptides were analysed on a timsTOF Pro 2 mass spectrometer in data-independent acquisition (DIA) mode, with parallel accumulation-serial fragmentation (dia-PASEF) and optimized scan windows (py_diAID, Ref.^91^) in a 350-1100m/z precursor scan range and ion mobility range of 0.63-1.45 1/K0, excluding single charged ions. Data acquisition was performed using Compass Data Analysis (v6.3.1.8).

### LC-MS/MS post-processing and Analysis

Precursor identification and quantification was done in Spectronaut (v19.5.241126.62635) against the Uni-Prot reviewed mouse reference proteome (December 2022) in library-free, specific Trypsin/P digestion mode with min. peptide length of 7 amino acids and max. 2 missed cleavages. Carbamidomethylation of cysteine residues was set as a fixed modification, while protein N-terminal acetylation and methionine oxidation were set as variable modifications. MaxLFQ was used for label-free quantification of protein groups. False discovery rate (FDR) threshold of precursor spectral matches, peptides and protein groups were set to 1%. Precursor mass deviations of up to 4.5 ppm and fragment mass deviations of 20 ppm were permitted. Subsequent bioinformatic analyses performed with Perseus (v1.6.15, Ref.^92^) and Python (v3.12). Before statistical analysis, quantified proteins were filtered for at least 60% valid values in at least one experimental group of replicates, and missing values were replaced sample-wise by random drawing of numbers from 1.8 standard deviations downshifted, and 0.3 width relative to the median and standard deviation of the distribution of valid values. Differentially expressed proteins were determined as Adj. *P* < 0.05, Welch’s T-test, Permutation based FDR with 250 randomisations.

### Quantification and Statistical Analysis

Sample sizes were based on similar previous studies. Mice were randomly allocated into treatment groups. Experimenters were not blinded to experimental groups during EAE scoring for synthetic diet fed mice. Experimenters were blinded to experimental groups during butyrate and propionate supplementation experiments. Experimenters were blinded for image analysis. Data are reported as mean ± s.e.m or s.d as indicated. With the exception of LC-MS/MS data, statistical analyses were performed using Prism v.10 (GraphPad). The Shapiro-Wilk test was used to assess normality of distribution, and Levene’s test was used to assess equality of variances, to determine which statistical test was appropriate for each comparison. For comparisons of two independent groups, an unpaired two-sided Student’s t-test was used when data were normally distributed and variances were equal; Welch’s t-test (unequal-variance t-test) was used when data were normally distributed but variances were unequal; and the Mann-Whitney U test was used when data were not normally distributed. For comparisons across three or more independent groups, the Kruskal-Wallis test with Dunn’s multiple comparisons post hoc test was used. For repeated-measures comparisons across three or more paired conditions, repeated-measures ANOVA with Holm-Šídák’s post hoc test was used. All specific tests as well as the number of biological replicates are described in each figure legend.

## Acknowledgements

We would like to thank Prof. Alexander Scheffold (Technische Universität Berlin) for providing input to the manuscript. We would like to thank the Flow Cytometry Core Facility and the Microscopy Core Facillity of the Medical Faculty at the University of Bonn for providing support and instrumentation funded by the Deutsche Forschungsgemeinschaft (DFG, German Research Foundation) – Project # 216372545, Project # 387333827, Project # 471514137 and Project # 388168919. We would like to thank the NGS Core Facility of the Medical Faculty at the University of Bonn/WGGC-Bonn for providing support and instrumentation for the scRNAseq and scTCRseq analysis. This work was funded by the Deutsche Forschungsgemeinschaft (DFG, German Research Foundation) under Germany’s Excellence Strategy – EXC2151 – 390873048 (to E.L. and F.M.), by the DFG SFB1454 - 432325352 (to E.L. and F.M.), by the state of Berlin and the European Regional Development Fund through the grant EFRE PersMedLab (to M.-F.M.), the Leibniz Association through the Leibniz Collaborative Excellence ImpACt project (to M.F.M. and A.K.), the German Center for Child and Adolescent Health (DZKJ) grant 01GL2401C (to M.-F.M.), the Faculty of Medicine at the University of Bonn BONFOR 1B Postdoc Fellowship Project ID: O-187.0008.1 (to F.D.W), and Leibniz Junior Research Grant J219/2025 (to F.D.W).

## Author Contributions

A.D, F.D.W and F.M conceived the study. F.D.W and F.M supervised the work. A.D and F.D.W designed experiments. A.D, B.G, A.P, S.R, T.S, Y.A, R.K and M.C performed experiments. A.D, B.G, J.B.K, A.M, A.P, S.R, F.H, P.D and F.R performed data analysis and visualisation. F.D.W, F.M and E.L acquired funding. F.D.W wrote the manuscript, with contributions from A.D and F.M. All authors reviewed and edited the manuscript.

